# Cut-Detector: A Tool for Automated Temporal Analysis of Late Cytokinetic Events

**DOI:** 10.1101/2025.06.06.658046

**Authors:** Thomas Bonte, Lucas Dubois, Paul Gagna, Rayane Dibsy, Anđela Petrović, Tamara Advedissian, Murielle Serres, Frédérique Cuvelier, Marie Crouigneau, Nathalie Sassoon, Neetu Gupta-Rossi, Stéphane Frémont, Arnaud Echard, Thomas Walter

## Abstract

Cytokinesis is the final step of cell division, resulting in the physical separation of the two daughter cells. Despite its fundamental importance in cell biology, biologists currently lack automatic tools for large-scale profiling of cytokinesis dynamics. In particular, the timing of the first microtubule cut in the intercellular bridge connecting the daughter cells is crucial, as it marks a critical step preceding abscission. Here, we introduce Cut-Detector, an open-source tool for the automatic analysis of late cytokinesis timing from time-lapse microscopy movies. Cut-Detector employs an AI approach to carry out the multiple tasks required to monitor cytokinesis: cell segmentation and tracking, detection of cell division events, localization of the intercellular bridge, and detection of the microtubule cuts. Cut-Detector will facilitate large-scale analyses to uncover new cytokinetic genes, a task that would be impractical without automation.

## Main

Cell division is an essential process in cell biology, which comprises two main phases: the segregation of the replicated chromosomes during mitosis followed by cytokinesis which leads to the formation of the two independent daughter cells. Cytokinesis eventually requires the cut of the intercellular bridge (ICB) that forms between the two daughter cells^1,2^. First, microtubules within the ICB are cut on one side of the midbody, an organelle within the ICB that orchestrates late cytokinetic events. Subsequently, microtubules on the opposite side of the midbody are cut, before the membrane cut (abscission) leading to the physical separation of the daughter cells^1,2^ (Online Methods, Extended Data Fig. 1).

Despite extensive research over several decades, our understanding of cytokinesis remains incomplete, in particular the late events that occur after furrow ingression^3–5^ (Extended Data Fig. 1). Traditionally, screens for identifying new cytokinetic genes rely on the detection of binucleated cells in fixed samples after selective gene inactivation using RNAi, which proved to be successful to identify new genes important for either furrow ingression or ICB stability^6,7^. However, upon inactivation, many important proteins in the final cytokinetic events do not lead to binucleated cells but rather a delay in abscission^8–12^. A key event preceding abscission is the first cut of the microtubules (MTs) within the ICB, that can only be followed by high-magnification time-lapse microscopy movies with a MT fluorescent marker^13–16^. However, quantifying the time of MT cut in such movies is tedious and requires extensive manual annotation by a trained biologist, and represents a bottleneck that precludes large-scale studies.

Here, we introduce Cut-Detector (Fig. 1a), a tool for the automatic analysis of the timing of the late cytokinetic events from time-lapse microscopy movies comprising phase contrast and two fluorescent channels highlighting microtubules (SiR-tubulin probe) and the midbody (GFP-MKLP1 fusion protein) (Extended Data Fig. 1). Cut-Detector employs an AI approach to carry out the tasks required to monitor cytokinesis: cell tracking (Fig. 1b, step 1), detection of cell division events (Fig. 1b, step 2), localization of the ICB (Fig. 1b, step 3) and detection of the microtubule cuts (Fig. 1b, step 4). As shown below, this allows the identification of proteins important for abscission by comparing the distributions of the MT cut timing in control vs. depleted cells.

**Fig. 1:**
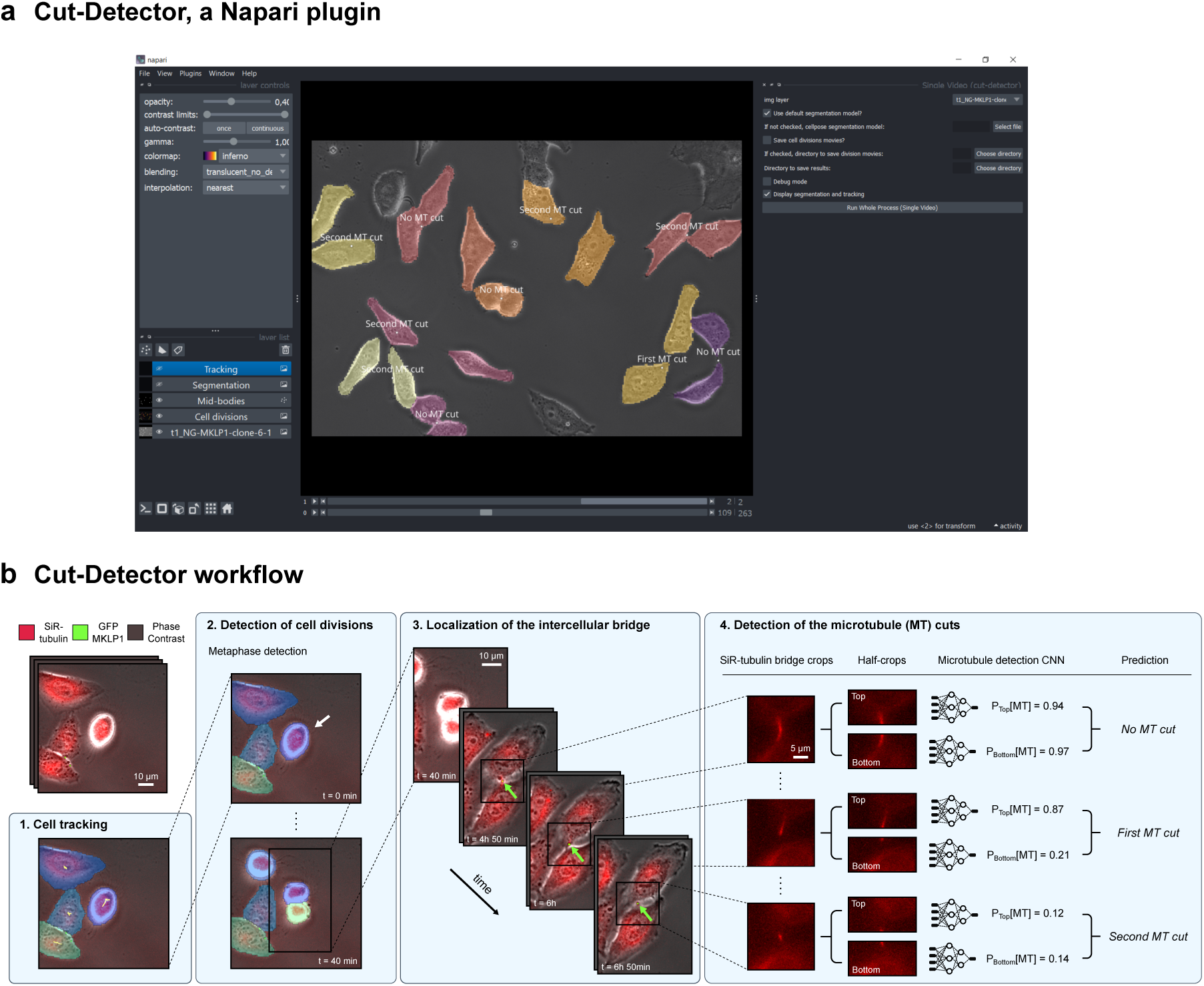
Step-by-step workflow of Cut-Detector and implementation as a user-friendly interface. **(a)** Cut-Detector has been developed within the Napari framework, enabling immediate user adoption. It allows cell tracking, cell segmentation, detected cell divisions, midbodies, and ICB classification to be visualized directly within Napari. **(b)** Overview of the Cut-Detector workflow. (1) First, cell segmentation is performed using a fine-tuned Cellpose segmentation model. Next, cell tracking generates cell tracks from segmented cell spots. (2) The classification of cells in metaphase (white arrow) allows for the identification of cell division events. (3) The midbody (green arrows), located at the center of the ICB, is detected in the GFP-MKLP1 channel. (4) Each SiR-tubulin crop is classified into one of the three classes: ICBs with no microtubule cut, ICBs after the first microtubule cut and ICBs after the second microtubule cut. The crop is divided into two half-crops on which a CNN model is applied to detect the presence of microtubules. The detection results from both halves are then combined to achieve accurate and robust classification.

The first microtubule cut defines a key step in cytokinesis and therefore needs to be detected with high accuracy and robustness. However, this is a difficult task to automatise given the low MT signal/noise (Extended Data Fig. 1, red channel). Here, Cut-Detector first extracts for each cell division time-lapse movie SiR-tubulin crops centered on the midbody detected by the GFP-MKLP1 marker. We then approach microtubule cut detection as a classification problem, defining 3 classes of ICBs: 1) without microtubule cut, 2) after the first microtubule cut, and 3) after the second microtubule cut (Fig. 1b, step 4). This classification task is challenging due to the brief interval between the first and second cuts, resulting in very few images between the two cuts in our dataset. Because of this class imbalance, a standard Convolutional Neural Network (CNN) classifier performs poorly (Online Methods).

Instead, we propose to detect the presence or absence of microtubules in each half of the image independently (Extended Data Fig. 2a). This approach leverages the fact that the ICB — whether microtubules have not been cut or after the microtubule cuts — spans across the image, as images are centered on the midbody located at the center of the ICB. Hence, microtubules will not be confined to a single half. We renamed the three microtubule cut detection classes as follows: 1) ICBs with microtubules present in both halves of the image, 2) in only one half of the image, or 3) in neither half of the image. From each original SiR-tubulin crop, we generated two half-images, each labeled with the presence or absence of microtubules, to train a microtubule detection CNN (Extended Data Fig. 2b-c). This strategy generates a large number of examples for both microtubule presence and absence, effectively addressing the issue of class imbalance. During inference, we similarly split our SiR-tubulin crop into two half-images and compute the probability of microtubule presence for each half (Extended Data Fig. 2d). We then combined these two predictions to perform the final classification (Fig. 1b, step 4).

We next used Cut-Detector in control cells and in cells treated with siRNAs to deplete either ALIX, Spastin or CEP55 — three proteins known to participate in abscission by promoting the MT cut^5,17–20^ (Online Methods) — or MICAL1 — whose inhibition was shown previously to delay the timing of the cell membrane cut^10^ but whether it does so by regulating the MT cut is unknown. The distributions of the MT cut timings detected by Cut-Detector from 10-min interval time-lapse movies in ALIX, Spastin and CEP55 depleted cells are significantly delayed (p-value < 0.05) compared to the control across three independent experiments — ALIX depletion being the only condition with one experiment —, as shown by Mann-Whitney U tests (Fig. 2a), as expected. Conversely, the distributions in MICAL1-depleted cells and in control cells were statistically equivalent within a margin of at most 30 minutes, as determined by a two-sided Mann-Whitney U test (Fig. 2a). Thus, MICAL1 is the first protein to be shown to promote cytokinetic abscission without modifying the timing of the MT cut.

**Fig. 2:**
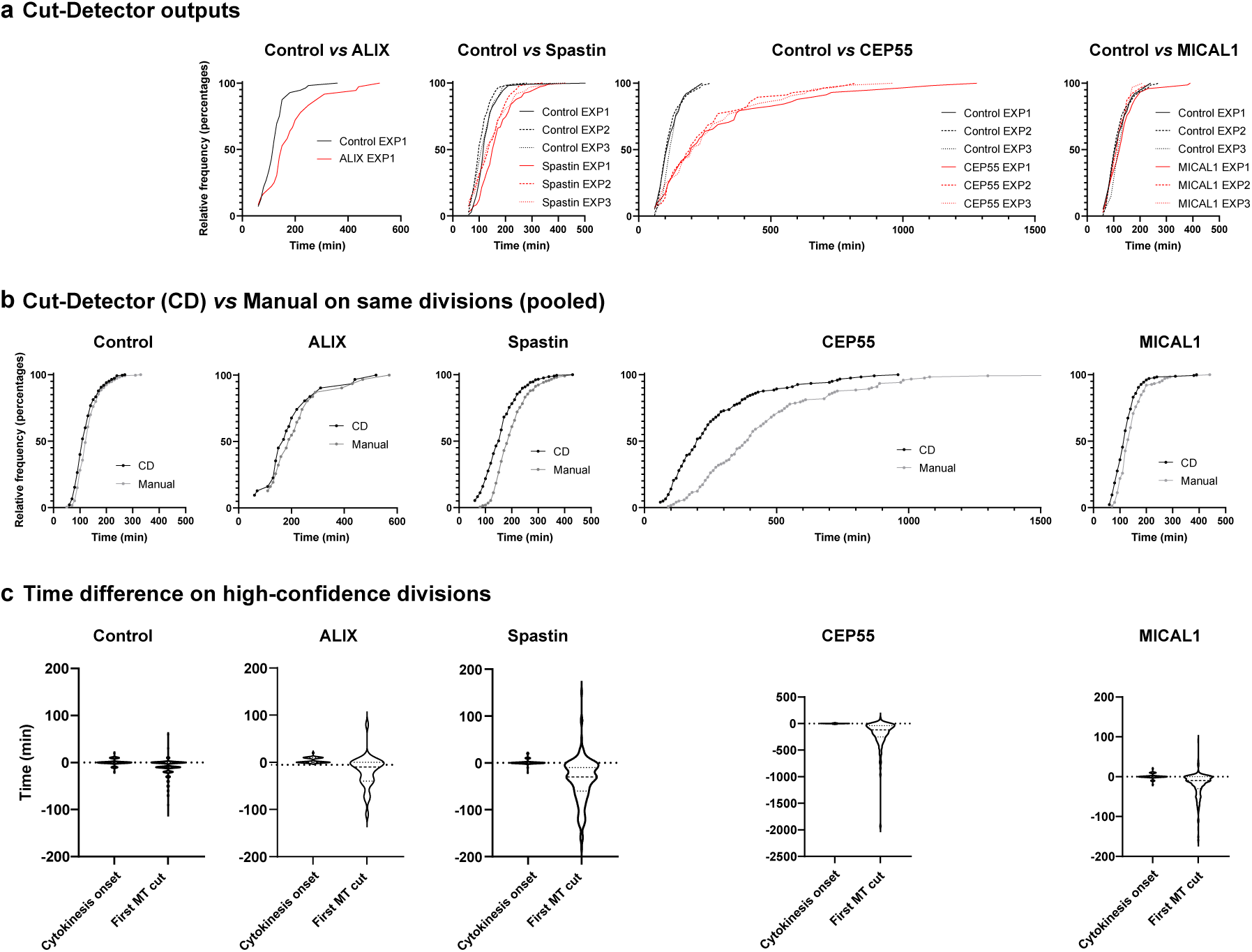
Performance of Cut-Detector in control cells and in cells depleted of ALIX, Spastin, CEP55 and MICAL1. Here and in the rest of the manuscript, “Relative frequency (%)” represents “Cumulative % of cells that completed the first MT cut” as a function of the time after furrow ingression. **(a)** Cut-Detector output: distributions of microtubule cut timings under conditions following either ALIX, Spastin, CEP55 or MICAL1 depletion, compared to a control distribution. Distributions for ALIX, Spastin and CEP55 are statistically different from the control, while MICAL1 showed statistical equivalence within a margin of at most 30 minutes. **(b)** Distributions of microtubule cut timings for divisions analyzed both by Cut-Detector and manually identified. Cut-Detector recapitulates the overall tendencies captured by manual annotation, albeit with statistical difference for Spastin and CEP55 depleted cells, putatively due to slow depolymerization of the MT whithin the ICB. The distributions of Cut-Detector vs. manual in control, ALIX and MICAL1 depleted cells are statistically equivalent within a margin of at most 40 minutes. **(c)** Detected time difference between Cut-Detector and manual annotations, for high-confidence divisions analyzed by both methods. Cytokinesis onset t_0_ and first microtubule cut t_1_ are analyzed separately.

To assess the accuracy of Cut-Detector, we compared its performance to manual annotations of the same movies. Fig. 2b shows the microtubule cut timing distributions for divisions detected by Cut-Detector and manually identified. The distributions of Cut-Detector vs. manual in the control, ALIX and MICAL1 depleted cells are statistically equivalent within a margin of 20, 40 min and 30 minutes, respectively, based on a two-sided Wilcoxon signed-rank test (Fig. 2b). However, the distributions of Cut-Detector vs. manual in the Spastin and CEP55 depleted cells are statistically different (Fig. 2b). This discrepancy can be explained because Spastin and CEP55 depletions prevent a clear microtubule cut and instead lead to gradual microtubule depolymerization – a process that is substantially longer and more challenging to detect accurately, even for a trained biologist (Extended Data Fig. 3, Online Methods). Of note, across all conditions, differences arise essentially from the detection of the MT cut time (t_1_), whereas the onset of cytokinesis (t_0_) is consistently well identified (Fig. 2c, Online Methods and Extended Data Fig. 1). Even for these challenging cases, Cut-Detector identified a significant delay in the MT cut in the depleted cells. Of note, Cut-Detector was able to detect, again, the absence (MICAL1 depletion) and the presence (ALIX, Spastin and CEP55 depletions) of MT cut delays in another cell line (expressing MgcRacGAP-mNeonGreen as a different midbody marker) that had not been used for training, showing that Cut-Detector can likely perform in several cell models (Extended Data Fig. 4, Online Methods).

We implemented Cut-Detector within the Napari framework as an open-source tool. As such, Cut-Detector enables immediate user adoption and allows for the visualization of cell tracking, cell segmentation, detected cell divisions, midbodies, and ICB classification directly. This establishes Cut-Detector as a valuable tool for quantifying the dynamics of late cytokinetic events. Cut-Detector will facilitate the identification of new genes that play a significant role in cytokinesis using high-content screening approaches, an important goal since cytokinetic failure is known to promote tumor formation in vivo^21^.

## Online Methods

### Cytokinesis

Cytokinesis is the last step of cell division and a multistep process that leads to the physical division of the daughter cells (Extended Data Fig. 1). Cytokinesis is essential for cell proliferation, tissue growth and renewal, and its dysfunction can lead to genetic instability and tumor formation^1,21^. It begins during anaphase with the formation of a cleavage furrow, whose invagination is driven by a contractile ring composed of actin and myosin filaments (Extended Data Fig. 1c-d). As contraction completes, the two daughter cells remain connected by an ICB (Extended Data Fig. 1e), at the center of which lies the midbody – a structure that concentrates key components required for abscission^1–2^. The MKLP1 kinesin has been shown to localize CEP55 to the midbody, where both proteins localize extensively. CEP55 then enables the recruitment of the Endosomal Sorting Complexes Required for Transport (ESCRT) machinery and associated proteins like ALIX which are responsible for abscission^5,17–19^. To do so, the ESCRT-III component CHMP1B first recruits the microtubule-severing enzyme Spastin which cuts the ICB microtubules on both sides of the midbody^9,20,22^. This step is a prerequisite for the membrane cut or abscission close to the site of the MT cut^15^. In parallel, clearance of the actin cytoskeleton from the ICB is crucial for abscission, and relies in part on the actin oxidase MICAL1^1,2,8–10,23–25^. However, proteomic analysis suggests that many proteins required for late cytokinetic events remain to be discovered^4,5,26^.

### Cell tracking

First, cell segmentation was performed using Cellpose^27^ on the phase contrast channel. We used the Cellpose interface to conduct human-in-the-loop fine-tuning of the default Cellpose segmentation model. This involved using 95 frames randomly selected from 53 time-lapse videos, including 50 that were outside our dataset and 3 in our train or validation sets. We retained Cellpose’s default training parameters: a learning rate of 0.1, weight decay of 0.0001 and 100 epochs. Segmentation was performed with both the flow threshold and cell probability threshold set to 0. The cell diameter was automatically calculated by Cellpose and set to 164 px, i.e. 36.9 μm. While Cut-Detector includes a default segmentation model, users can easily switch to any model fine-tuned from Cellpose, facilitating adaptation to slightly different data and cell lines.

Although cell properties such as cell division can be used for tracking, the high temporal resolution of our data allowed us to rely solely on particle tracking strategies. Particle tracking typically focuses on tracking spot points, but it can be adapted for cells by using the barycenter of cell segmentation masks. We used the particle tracking algorithm implemented in LapTrack^28–30^. Such tracking involved matching cell spots between consecutive frames. The core idea is to define a cost function that estimates how likely two cell spots belong to the same cell track. A smaller cost indicates a higher probability that the spots belong to the same track. This cost function uses features of the spots (like circularity or signal intensity), which can be selected based on the specific application. Tracking involved pairing spots to minimize the total cost function.Given two spots A and B, the cost function C between these two spots is given by the following equations:

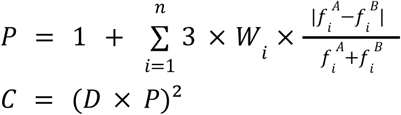

where W_i_ refers to the weight associated with feature f_i_ and D is the Euclidean distance.

Here, the high time resolution allowed us to rely only on the distance between cells, with no additional features used (P = 1). The maximum distances for tracking and gap closing was set to the Cellpose’s cell diameter and half of the Cellpose’s cell diameter, respectively. Gap closing was allowed for up to three frames. Track merging and splitting were not permitted. Tracks with fewer than 10 spots were ignored, as they were likely to result from incorrect segmentation.

### Detection of cell division events

Detection of cell division relies on identifying metaphase cells, which is approached as a classification problem. Metaphase cells adopt a distinctive round shape, making them relatively straightforward to identify (Extended Data Fig. 1b).

We trained a CNN classification model to recognize metaphase cells in cell tracks. Our classifier was trained and evaluated on 12 time-lapse movies, including 9 outside our main dataset: 8 time-lapse videos for training, 2 for validation, and 2 for testing. We applied Cellpose segmentation to these 12 time-lapse videos and extracted crop images of each detected cell at each frame. These images were then manually annotated into one of three classes: interphase cell, metaphase cell or dead cell. We ended up with a total of 3,257 annotated images of size 256×256: 1,217 interphase cells, 1,192 metaphase cells, and 848 dead cells, with all three channels (phase contrast, SiR-tubulin, GFP-MKLP1).

For this classification task, we used a ResNet50 model pre-trained on ImageNet and data augmentation such as rotation, horizontal flip, and vertical flip. We used an Adam optimizer to minimize the cross-entropy loss. The model was trained for 500 epochs with a learning rate of 5e-5, taking six hours on a GPU. The accuracy on the test set was 0.99.

We used Hidden Markov Models (HMM) at inference to correct for potential inconsistencies, such as detecting interphase cells after cell death within the same cell track^31^. HMM parameters were determined using the Baum-Welch algorithm on 200 cell tracks extracted from our training time-lapse videos, setting the probability of transition from death to interphase or metaphase to 0. Corrections were applied using the Viterbi algorithm on the ResNet50 predictions. Metaphase events were ultimately defined as sequences of consecutive metaphase spots.

To detect cell division events, we iterated over all cell tracks starting at frame>0, i.e. likely to be daughter tracks. For each daughter cell track, we searched for metaphase events in its vicinity. If multiple candidates were found, we ranked them based on their Intersection over Union (IoU) over the first daughter cell frame. Finally, we merged the mother cell track associated with the metaphase event to the daughter cell track, defining a cell division event. The onset of cytokinesis, t_0_, is defined at the first frame of the daughter cell track and corresponds to the time of furrow ingression

The four time-lapse videos of our test set contain 96 cell divisions, including abnormal tripolar divisions. 86 divisions were correctly detected, while three events were identified that did not correspond to any actual cell division (recall 0.90, precision 0.97).

### Localization of ICBs

Since SiR-tubulin images can be noisy (see examples in Extended Data Fig. 3, red channel), locating the ICB only from the SiR-tubulin channel is particularly challenging. To address this issue, we used the GFP-MKLP1 channel to detect the midbody, which is located at the center of the ICB and is highly enriched for MKLP1. Midbody detection is performed in three steps, independently for each cell division. First, midbodies are detected within the cell division crops. It is important to note that even within a single cell division, there may be multiple midbodies, either from closely positioned ICBs or remnants associated with cells from previous divisions^32^. Second, these detected midbody spots are used to generate midbody tracks over time. Finally, the track associated with the specific cell division is selected.

Since midbodies appear as blobs in the GFP-MKLP1 channel images, we used the widely adopted difference-of-Gaussians method for their detection. Specifically, we utilized the difference-of-Gaussians function from the skimage package^33^, with sigma going from 2 to 5 with a ratio of 1.2, and a threshold set to 0.1. These parameters were chosen to correspond to the typical size of midbodies in GFP-MKLP1 images (typically 1 μm). The images were normalized before applying the difference-of-Gaussians function.

For midbody tracking, we used the same framework as for cell tracking^28–30^. We incorporated two additional penalties: intensity in the SiR-tubulin channel with a weight of 1.5, and intensity in the GFP-MKLP1 channel with a weight of 5. The maximum distance for tracking and gap closing was set to 175 px, i.e. 39.375 μm. Gap closing was permitted for up to three frames, while track merging and splitting were not allowed. If multiple midbody tracks were detected, we first eliminated those with fewer than five frames and whose average SiR-tubulin intensity was below the third quartile of the image within the first 100 minutes of cytokinesis. This is because the correct midbody is located roughly at the center of the ICB, and thus exhibits a significant SiR-tubulin signal. The remaining candidates were then sorted based on their distance to the barycenter of the daughter cells, which is the expected position of the midbody. We finally selected the closest candidate.

Midbody detection is considered accurate if the midbody position was within 10 px of the correct location for at least 90% of the frames between the onset of cytokinesis t_0_ and the second microtubule cut t_2_ (Extended Data Fig. 1f-g). Since no training was required, we tested our method on the entire dataset, yielding a midbody detection accuracy of 0.91. Most of the missed midbody tracks were due to a fading GFP-MKLP1 signal after the first microtubule cut, which made detection significantly more challenging, and in some cases, impossible.

### Detection of the microtubule cut

The detection of the microtubule cut was approached as a classification problem. We extracted 6,878 SiR-tubulin ICB crops from 199 cell divisions in our dataset. The crops were 100×100 px images centered on the annotated midbody. They were manually labeled by a trained biologist into three classes: 2,929 with no microtubule cut, 355 after the first microtubule cut, and 3,594 after the second microtubule cut. Although our primary interest is the first microtubule cut, we distinguished between one and two cuts for further analysis.

We first trained a ResNet18 classifier to perform such classification. Data augmentations included rotation, horizontal flip, and vertical flip. Images were normalized before training. We used an Adam optimizer to minimize the cross-entropy loss. The model was trained for 100 epochs with a learning rate of 1e-4, taking 50 minutes on a NVIDIA RTX A3000 Laptop GPU. Early stopping was employed to select the best model based on validation performance.

To account for class imbalance, the classification performance was evaluated using macro-averaged accuracy. Macro-averaged accuracy was computed by averaging the classification accuracy across all classes, giving equal weight to each class. Our model achieved a score of 0.73 on the test set. This relatively low performance can be attributed to the significant class imbalance, which hampered the model’s ability to accurately classify images between the first and second microtubule cuts.

Since ICB images are centered on the midbody, the ICB spans across the image and microtubules are not confined to a single half. This led us to adopt a different approach, reframing the problem as detecting the presence or absence of microtubules in each half of the image independently. Accordingly, we renamed the three classes of ICBs with no MT cut, after the first MT cut and after the second MT cut as: ICB with microtubules present in both halves of the image, present in only one half of the image and present in neither half of the image, respectively.

To enhance the robustness of our approach, we introduced four distinct ways of splitting images into half: along the horizontal axis, the vertical axis, and the two diagonals (Extended Data Fig. 2a). Using this strategy, we generated eight half-images per original image: top, bottom, left, right, top-left, bottom-right, bottom-left and top-right. Each half-image was labeled with the presence or absence of microtubules, to train a microtubule detection CNN. If the original image corresponded to the stage after the second microtubule cut, then all eight half-images were labeled with the absence of microtubules. Otherwise, the exact positions of microtubules had to be identified to assign correct labels. We measured SiR-tubulin intensity along a circular profile of radius 11 px, i.e. 2.5 μm, centered on the midbody (Extended Data Fig. 2b). This value was chosen as microtubules are typically cut at 1-2 μm from the midbody. We used the find_peaks function from the scipy package^34^ to identify the one or two local maxima corresponding to the microtubules. These positions allowed us to label our half-images into containing microtubules, or not. If the axis of symmetry overlapped with detected microtubules, the corresponding half-image pair was excluded to avoid ambiguous labeling. It is important to note that this microtubule detection method effectively labeled half-images because it leverages prior knowledge of how many microtubule cuts occurred in the original image, which was manually annotated by a trained biologist. This allowed us to determine precisely how many local maxima to identify along the circular profile. In contrast, when applied directly as a classification tool, this method failed to generalize to unseen data, performing comparably to the ResNet18 model.

Finally, we applied mirroring to the half-images (Extended Data Fig. 2c) to make them square. This mirroring approach not only allowed us to use standard CNN models but also provided immediate data augmentation within the images themselves.This process resulted in a synthetic dataset with a total of 45,176 images of size 100×100 px, including 15,368 labeled with microtubules and 29,808 labeled without.

We next trained a ResNet18 model, pretrained on ImageNet, for this microtubule detection task. Data augmentations included rotation, horizontal flip, and vertical flip. Images were normalized before training. We used an Adam optimizer to minimize the cross-entropy loss. The model was trained for 100 epochs with a learning rate of 1e-4, taking height hours on a NVIDIA Tesla P100 GPU. Early stopping was employed to select the best model based on validation performance. The accuracy on the synthetic test set is 0.95.

During inference, each image is processed to generate the eight mirrored half-images (Extended Data Fig. 2d). We computed the probability P[MT] of microtubule presence independently for each half-image, using the microtubule detection CNN. These probabilities were then combined pair-wise as follows: top/bottom, left/right, top-left/bottom-right, and bottom-left/top-right. This pairing allowed us to estimate the probability that the original image contains microtubules in both halves, in only one half, or in neither half. These corresponded to the three classification categories: no microtubule cut, after the first microtubule cut, and after the second microtubule cut. For example, for the top/bottom pair:

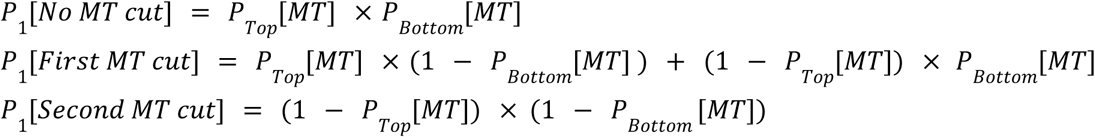

The same procedure was applied to the other three pairs of half-images to obtain P_2_, P_3_ and P_4_. To enhance robustness, these probabilities were averaged to get the final classification probabilities. This strategy effectively handled cases where microtubules aligned with a specific axis. For example, in the left/right split shown in Extended Data Fig. 2d, the microtubule detection model failed to detect microtubules in either half due to ICB alignment along the vertical axis. Averaging across multiple splits mitigated such misclassifications by incorporating information from alternative orientations.

The macro-averaged classification accuracy on the test set was 0.80, thus clearly outperforming direct classification (0.73). Extended Data Fig. 3 displays examples of ICB images, including wrong classifications. The errors were primarily attributed to highly noisy images, microtubules cutting near the edge of the image, or the presence of bright areas that significantly complicate cut detection.

Finally, similar to metaphase classification, we used Hidden Markov Models (HMM) to correct inconsistencies within a given cell division. Here, we set the probability of several transitions to 0: from one cut to zero cut, from two cuts to one cut and from two cuts to zero cut.

The first microtubule cut time, t_1_, was then identified at the first frame classified as “First MT cut”. In our test set, the difference between the predicted and ground truth first cut frames was -0.4 ± 3.0 frames across 49 cell divisions. This discrepancy exceeded four frames in only 6 of these 49 divisions.

### Selection of high-confidence mitotic events

Cut-Detector applied a series of algorithmic steps (cell segmentation, cell tracking, metaphase detection, midbody detection, midbody tracking, microtubule cut detection), each achieving high, yet not perfect, accuracy. Moreover, the performance of these steps varied depending on experimental conditions. For example, segmentation was highly accurate in low-cell-density scenarios but tended to be less reliable when cell density increased, particularly toward the end of experiments. Since these errors accumulate, it is crucial to identify cell divisions where measurements can be considered highly reliable.

To address this, Cut-Detector performed a series of quality control checks to focus exclusively on high-confidence mitotic events. Specifically, it excluded divisions in which the midbody was not detected or in which no cut was detected, tripolar divisions, and divisions where the midbody is located within 4.5 μm of the field-of-view edge. Furthermore, divisions where the microtubule cut was detected within 60 minutes of cytokinesis onset are disregarded, as such events are highly improbable and account for less than 0.5% of manually annotated divisions (11 out 2332). Supplementary Table 1 summarizes both the Cut-Detector and ground truth categories for all divisions from the four videos in our test set.

These quality control filters reduced the number of analyzed divisions compared to a human annotator. Specifically, Cut-Detector did not analyze 59.0%, 81.2%, 66.1%, 60.8%, and 70.2% of control, ALIX, Spastin, CEP55 and MICAL1 divisions, respectively. To obtain these values, we performed an automatic matching in which a Cut-Detector division was paired with its ground truth counterpart if the detected cytokinesis onsets differed by no more than 20 minutes, and the associated midbody positions were within a maximum distance of 125 px, i.e., 28 μm.

To ensure that missing divisions did not introduce any bias in microtubule cut detection, we compared the distributions of the ground truth microtubule cut times t_1_-t_0_ between divisions analyzed by Cut-Detector and those missed by Cut-Detector.

These distributions are presented in Extended Data Fig. 5, separately for the three experiments and five depletion conditions: control, ALIX, Spastin, CEP55 and MICAL1. We performed two one-sided Mann-Whitney U tests to assess the statistical equivalence of independent distributions. For all experiments in the control, ALIX, Spastin and MICAL1 conditions, the distribution of ground truth microtubule cut times for divisions detected by Cut-Detector and the distribution for divisions missed by Cut-Detector were statistically equivalent within a margin of at most 20, 40, 40 and 40 minutes, respectively. CEP55 depletion was the only condition where all tests were not significant at a margin of 40 minutes, which can be partially attributed to the much larger average cut time compared to other conditions. Nevertheless, we observe in Extended Data Fig. 5 that CEP55 distributions were similar to each other. This reinforces the conclusion that no bias in microtubule cut detection was introduced by the quality control filters of Cut-Detector.

Conversely, some events were identified by Cut-Detector as cell divisions, but no corresponding matches were found in the divisions annotated by biologists. These cases include divisions missed by annotators, divisions missed by our matching process, or false positive events. While the latter could potentially introduce bias in microtubule cut detection, in our first experiment they account for only 4.24%, 5.56%, 7.55%, 10.34% and 4.05% for control, ALIX, Spastin, CEP55 and MICAL1, respectively, making their impact on cut detection timing negligible.

### Independent evaluation of cytokinesis onset and first microtubule cut detections

The primary variable of interest in this study is the time interval between the onset of cytokinesis and the first microtubule cut, i.e. t_1_-t_0_. Consequently, it is essential to independently evaluate the detection of both the onset of cytokinesis (t_0_) and the first microtubule cut (t_1_). Fig. 2c illustrates the time detection differences for these two values, with data from the independent experiments pooled together. To assess the statistical equivalence of the paired distributions, we conducted two one-sided Wilcoxon signed-rank tests.

The distribution of the true onset of cytokinesis times and those detected by Cut-Detector were statistically equivalent within a margin of 10 minutes for all conditions. Similarly, the distribution of the true first microtubule cut times and those detected by Cut-Detector were statistically equivalent for the control, ALIX and MICAL1 depletion conditions within a margin of 20, 30 and 20 minutes, respectively. However, for Spastin and CEP55 depletion conditions, the test was not significant at a margin of 40 minutes, aligning with the observed difficulty in predicting the cut time for these conditions, as shown in Fig. 2b. These results confirmed that the main challenge lies in detecting the first microtubule cut time, whereas the detection of cytokinesis onset is achieved with high accuracy.

### Evaluation on a different cell line using a different midbody marker

To assess the robustness of Cut-Detector, we tested it on a different HeLa cell line that stably expressed MgcRacGAP-mNeonGreen instead of GFP-MKLP1 as the midbody marker. We performed one experiment for each of the previously described conditions.

Focusing on divisions detected both by Cut-Detector and manual annotators, we found that Cut-Detector detected the microtubule cut timing with comparable accuracy to that observed in the initial cell line (see Extended Data Fig. 4b-c). The respective distributions were statistically equivalent within a margin of 10 minutes for control, ALIX and MICAL1 depletion conditions, and within 40 minutes for the Spastin depletion condition, as determined by a two-sided Wilcoxon signed-rank test. Once again, CEP55 was the only condition for which the distributions significantly differed, consistent with the challenges previously observed in the initial cell line.

Interestingly, and as in the initial cell line, the MICAL1 depletion condition was the only depletion condition that showed statistical equivalence with the control, as determined by a two-sided Mann-Whitney U test and within a margin of 10 minutes. Conversely, ALIX, Spastin, and CEP55 depletion conditions were statistically different from the control (Extended Data Fig. 4a). This result indicates that MICAL1 depletion does not significantly impact microtubule cut timing, consistent with the observations from the GFP-MKLP1 cell line.

### Image data acquisition and annotation

HeLa cells were plated on glass bottom 12-well plates (MatTek) and incubated in an open chamber (Life Imaging) at 37 °C and equilibrated in 5% CO_2_. Time-lapse microscopy was performed by recording cells every 10 minutes over a 48-hour period using an inverted Nikon Eclipse TiE microscope equipped with CMOS MOMENT camera (Teledyne Photometrics) and a ×40 0.95 NA Plan APO λ objective, controlled by Metamorph software (Molecular Devices). Each pixel corresponds to 225 nm, with frame dimensions of 1600×1100 pixels. SiR-tubulin 5 nM (Spirochrome), a fluorescent probe that labels MTs, was incubated prior to imaging. In the analysis described above, the SiR-tubulin channel was a maximum projection of 4 z-stacks of 3 μm range.

Trained biologists manually annotated 13 time-lapse videos from four independent experiments, resulting in 199 cell divisions (Supplementary Table 2). We only considered divisions that started before the first 36 hours to ensure the entire division occurred within the duration of the video. All models were trained and tested on these 13 videos, except for cell segmentation and metaphase classification, which also utilized additional time-lapse videos since these tasks did not require annotated cell divisions. The entire analysis of the test data set by Cut-Detector took 31 minutes for four time-lapse videos, averaging 7 minutes and 52 seconds per video on a NVIDIA RTX A3000 Laptop GPU.

Additionally, Cut-Detector was evaluated on time-lapse videos acquired with control and depleted cells, as detailed below. To ensure reproducibility, we conducted three independent experiments for each condition, except for ALIX depletion, for which only one experiment was conducted.

### Cells and cell culture

HeLa cells stably expressing GFP-MKLP1 (generated as in Addi et al.^5^) and HeLa cells stably expressing MgcRacGAP-mNeonGreen (generated by the transposition of the plasmid coding for the fusion protein) were grown in Dulbecco’s Modified Eagle Medium (DMEM) GlutaMax (31966; Gibco, Invitrogen Life Technologies) supplemented with 10 % fetal bovine serum and penicillin–streptomycin (Gibco) at 50 ng/mL in 5 % CO_2_ at 37 °C.

### siRNA transfections

For silencing experiments, HeLa cell lines were transfected with 50 nM siRNAs (siControl, siSpastin, siCEP55, siMICAL1) or 20 nM (siALIX) for 5 days using Lipofectamine RNAiMAX (Invitrogen), following the manufacturer’s instructions.

siRNAs against Control non-targeting siRNA (5’UGGUUUACAUGUUGUGUGA[dU][dU]3′), ALIX (5’CCUGGAUAAUGAUGAAGGA[dU][dU]3′), Spastin (5’AAACGGACGUCUAUAAUGA[dU][dU]3′), CEP55 (5’UUUCUUAAGGAGCUCCGAAA[dU][dU]3′), and MICAL1 (5′GAGUCCACGUCUCCGAUUU[dU][dU]3′).

### Antibodies and plasmids

The following antibodies were used for Western blot experiments: rabbit anti-MICAL1 (1:500, Proteintech Europe 14818-1-AP), mouse anti-CEP55 (1:1000, Santa-Cruz - B-8 - #sc-374051), rabbit anti-Spastin (1:2000, ProteinTech - 22792-1-AP), mouse anti-ALIX (1:500, Biolegend – 634502) and mouse anti-GAPDH (1:10000, Proteintech, 60004-1-Ig).

### Western blot

Cells were lysed on ice for 30 min in Triton lysis buffer (50 mM Tris pH 8, 150 mM NaCl, 1 % Triton X-100 and protease inhibitors) and centrifuged at 10 000 g for 10 min. Proteins were separated in 4-12 % gradient SDS-PAGE or in 10 % SDS-PAGE (Bio-Rad Laboratories) and transferred onto PVDF membranes (Millipore). Membranes were blocked and incubated with indicated antibodies in 5 % milk in 50 mM Tris-HCl pH 8.0, 150 mM NaCl, 0.1 % Tween20, followed by horseradish peroxidase-coupled secondary antibodies (1:10,000, Jackson ImmunoResearch) and revealed by chemiluminescence (GE Healthcare) (Extended Data Fig. 6).

## Acknowledgements

This work was supported by the French government under management of Agence Nationale de la Recherche (ANR) as part of the “Investissements d’avenir” program (reference ANR-19-P3IA-0001; PRAIRIE 3IA Institute). Additionally, this work was supported by a government grant managed by the Agence Nationale de la Recherche under the France 2030 program, with the reference number ANR-24-EXCI-0004. This work was also supported by the Institut Pasteur, CNRS, FRM (Recherche soutenue par la FRM EQU202103012627) and the Agence Nationale pour la Recherche (ANR-20-CE011-0014 SepScort) to AE, as well as by Q-LIFE (ANR-17-CONV-0005) to AE and TW. RD was supported by Fondation ARC pour la recherche sur le cancer (POST-DOC2023080006931). TA received a post-doctoral fellowship from Fondation pour la Recherche Médicale (FRM SPF201809006907).

## Code availability

Cut-Detector can be downloaded from the Napari Hub: https://www.napari-hub.org/plugins/cut-detector. The Python code is available via GitHub: https://github.com/15bonte/cut-detector.

## Supplementary information

**Extended Data Fig. 1:**
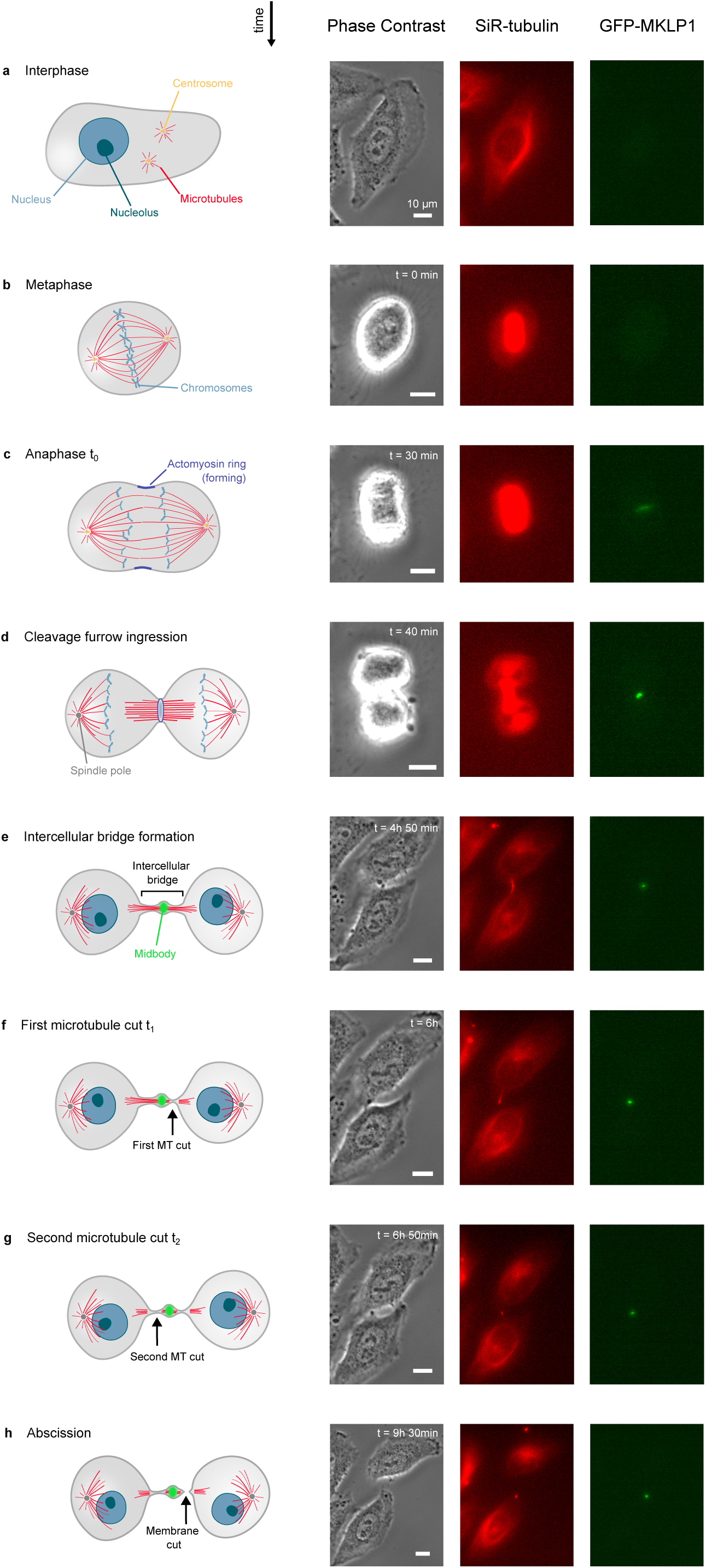
Cell division and the successive steps of cytokinesis. **Left panels:** schematic steps of cell division. **Right panels:** corresponding images from a representative movie of a HeLa cell stably expressing the midbody marker GFP-MKLP1 (green channel) and stained with the microtubule (MT) probe SiR-tubulin (red channel). The phase contrast channel is also provided. **(a)** During interphase, cells are flat and chromosomes are replicated before cell division. **(b)** In metaphase, mitotic cells have a characteristic round shape and align their chromosomes at the equator of the spindle, with centrosomes located at opposite poles. **(c)** In anaphase, the mother cell elongates and sister chromatids are separated. A contractile ring made of actin filaments and myosin II forms at the cell equator. This step marks the onset of cytokinesis (time t_0_). **(d)** The contractile ring drives complete cleavage furrow ingression. **(e)** At the end of contraction, the plasma membrane of the cleavage furrow narrows and the intercellular bridge (ICB) filled with MTs is present. The midbody is located at the center of the ICB. **(f)** Typically two hours after the cytokinesis onset, microtubules are then cut on one side of the midbody, typically at a distance of 1-2 µm, defining the first microtubule cut (time t_1_). **(g)** Microtubules are then cut on the other side, defining the second microtubule cut (time t_2_). **(h)** Finally, the cell membrane is cut (abscission), physically separating the two daughter cells.

**Extended Data Fig. 2:**
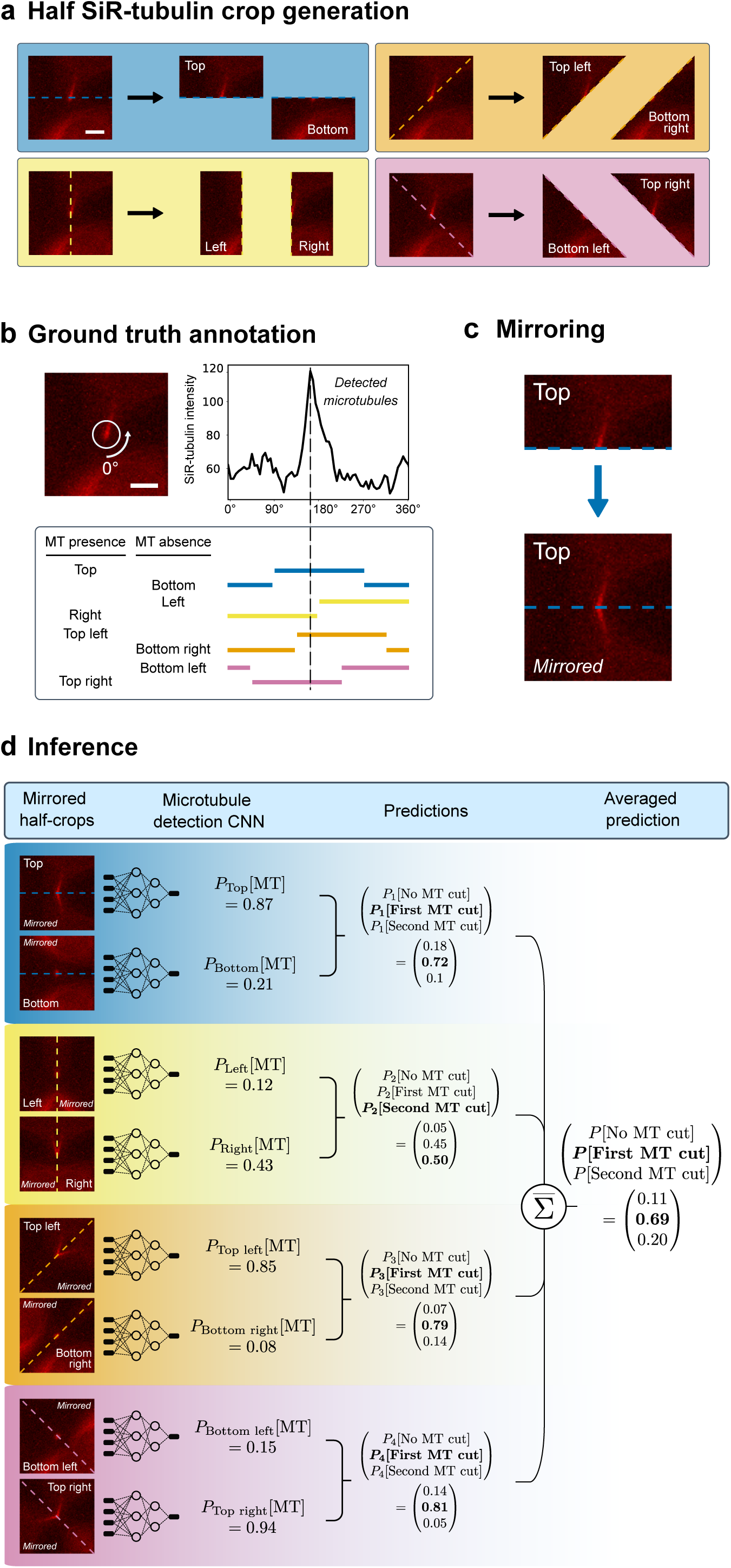
Methodology to define whether the ICB experienced zero, one or two MT cuts in a still image. **(a)** SiR-tubulin crops are split into half along four distinct axes: horizontal, vertical, and the two diagonals. Scale bar is 5 μm. **(b)** To train a microtubule detection CNN, each half-image was labeled with the presence or absence of microtubules. Here, since the original crop is taken after the first microtubule cut, the position of the remaining microtubules is determined by identifying the point of maximum intensity along a circular profile with a 2.5 µm radius. This position allows accurate labeling of each half-image based on whether it contains the remaining microtubules. Scale bar is 5 μm. **(c)** To fit standard CNN input requirements, half-images are mirrored to create square-shaped inputs. **(d)** During inference, the same half-images are generated as in (a). A trained microtubule detection CNN computes the probability of microtubule presence independently for each half-image. These probabilities are paired, and the results are averaged across the four pairs to improve classification robustness.

**Extended Data Fig. 3:**
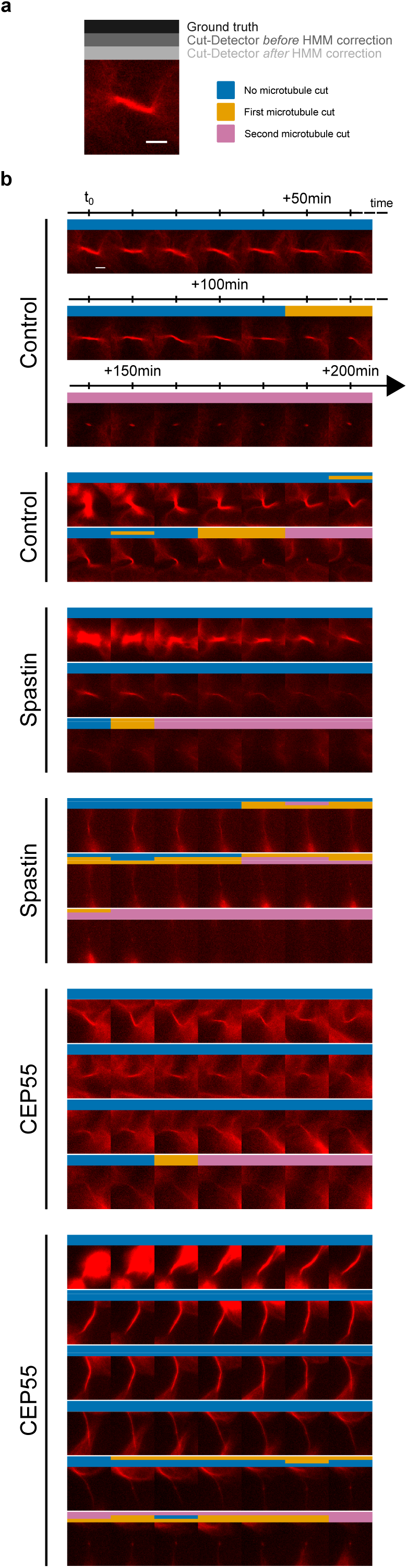
Challenges in detecting the microtubule cuts in depleted conditions when microtubules slowly depolymerize (Spastin and CEP55 depletions). SiR-tubulin intercellular bridge crops from movies in control cells and in cells depleted for either Spastin, or CEP55 (two representative movies per condition). **(a)** Each crop is annotated with the ground truth label and Cut-Detector predictions, both before and after applying a Hidden Markov Model (HMM), which helps resolve inconsistencies. Scale bar is 5 μm. **(b)** Time-lapse galleries of bridge crops, for the different conditions. Scale bar is 5 μm.

**Extended Data Fig. 4:**
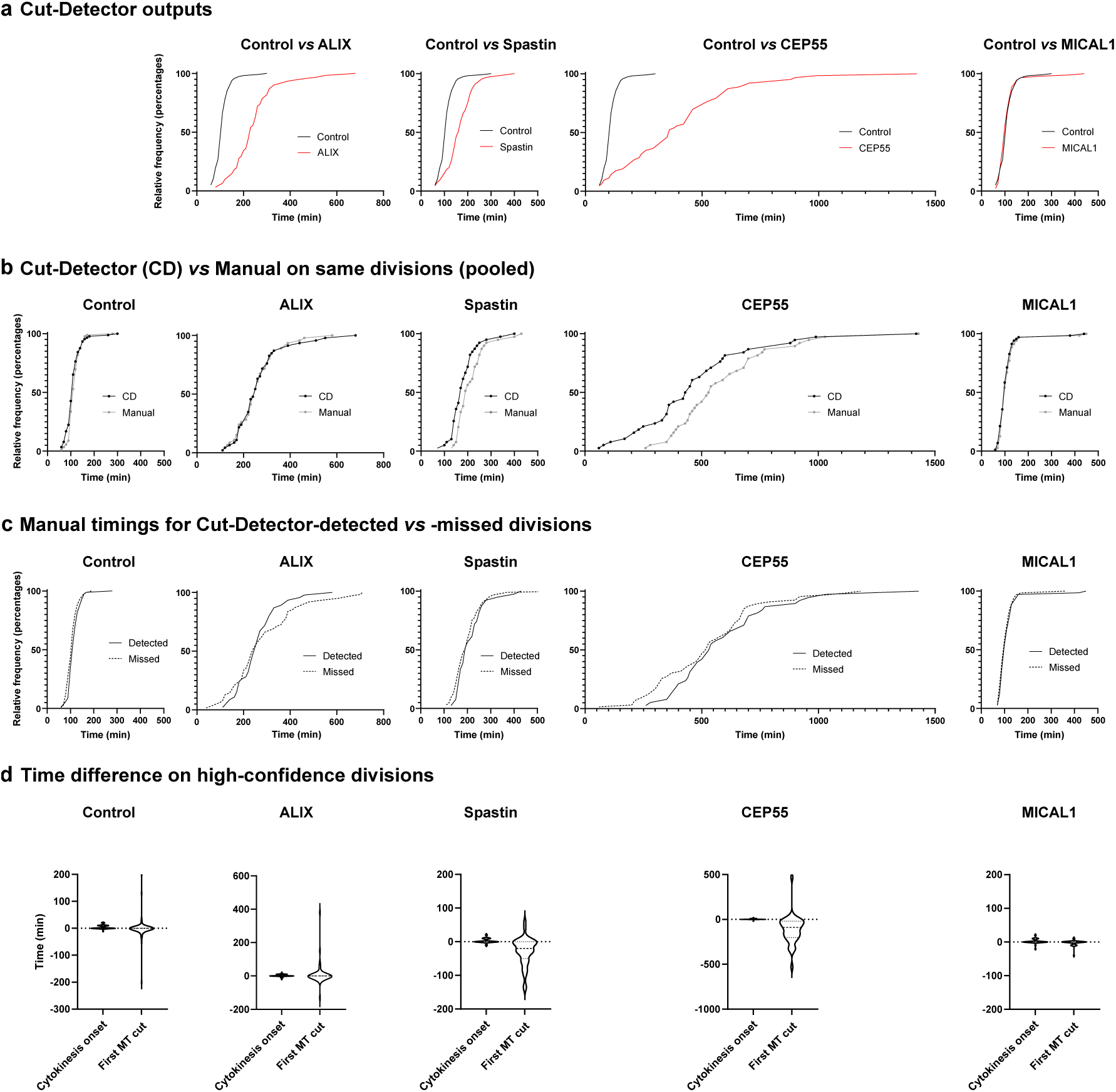
Performance of Cut-Detector in a MgcRacGAP-mNeonGreen HeLa cell line that has not been used for training. A HeLa cell line that stably expresses the midbody marker GFP-MgcRacGAP has been recorded by time-lapse microscopy in the presence of SiR-tubulin. **(a)** Comparaison of the distributions of microtubule cut timings detected by Cut-Detector in control cells versus cells depleted of either ALIX, Spastin, CEP55 or MICAL1. **(b)** Comparaison of the distributions of microtubule cut timings for divisions analyzed both by Cut-Detector and manually identified. **(c)** Comparaison of the distributions of manual microtubule cut timings for high-confidence divisions detected by Cut-Detector versus for divisions missed by Cut-Detector. **(d)** Detected time difference between Cut-Detector and manual annotations for high-confidence divisions analyzed by both methods. Cytokinesis onset t_0_ and first microtubule cut t_1_ are analyzed separately.

**Extended Data Fig. 5:**
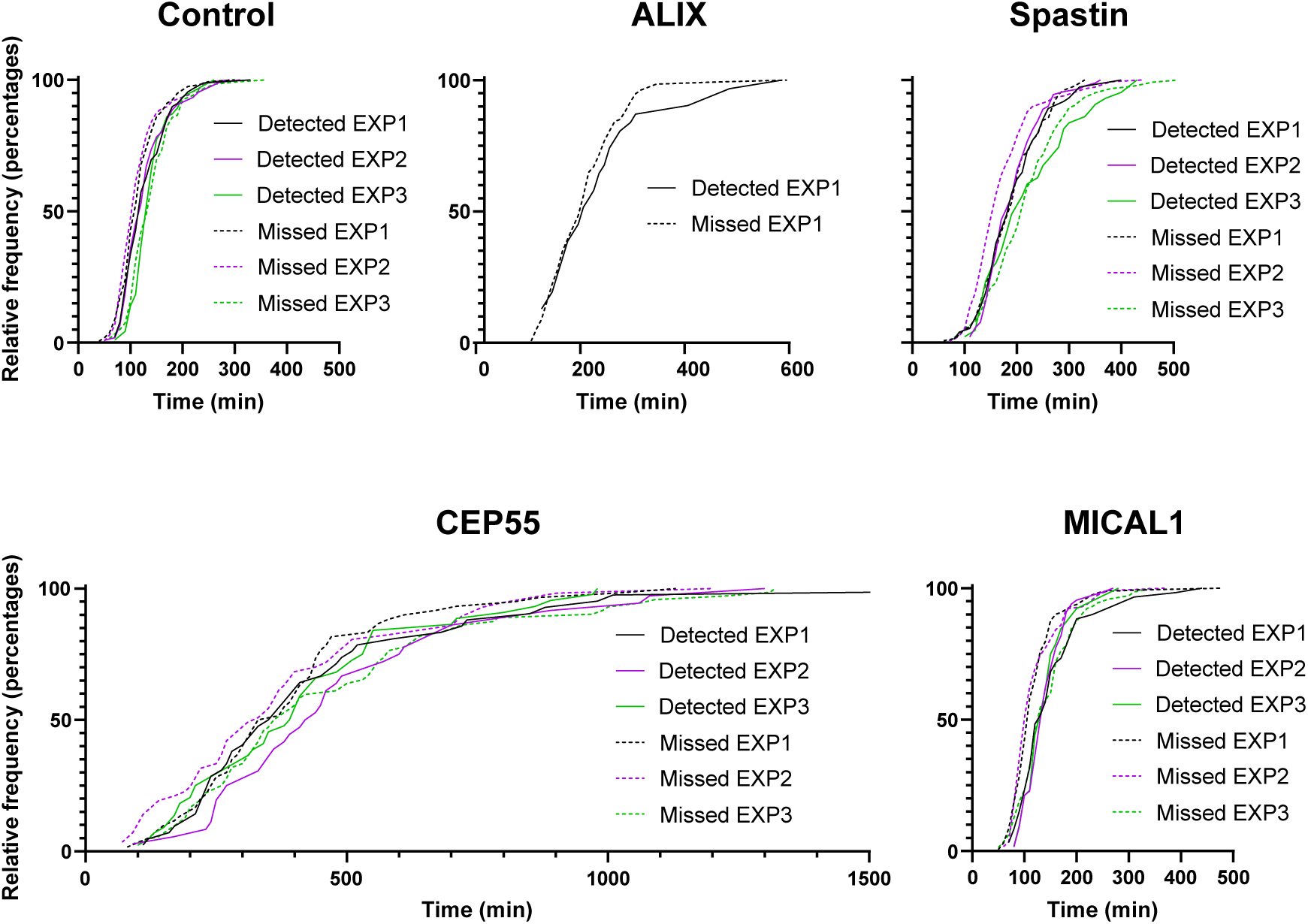
Comparison of manual microtubule cut timings for Cut-Detector-detected and -missed divisions. Distributions of manual microtubule cut timings for divisions automatically detected and non-detected, respectively. The corresponding distributions of microtubule cut times were statistically equivalent within a margin of at most 40 minutes for all experiments, with CEP55 being the only exception due to the high cut-time values. These results suggest that there is no bias introduced by the selective high-confidence cell division detection employed by Cut-Detector.

**Extended Data Fig. 6:**
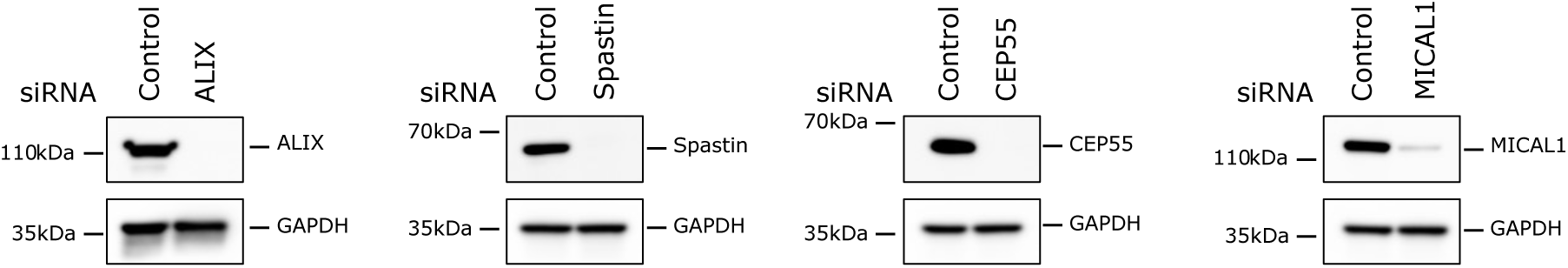
Western blot analysis showing efficient protein depletion. Western blot analysis of ALIX, Spastin, CEP55 and MICAL1 in GFP-MKLP1 HeLa cells treated with indicated siRNAs. Loading control: GAPDH. This experiment was repeated three times independently with similar results, for all conditions except ALIX.

**Supplementary Table 1:** Cut-Detector and ground truth categories of all divisions from the four videos in our test set. Notably, only 10 divisions out of 96 are missed by Cut-Detector, while 3 events were identified that did not correspond to actual cell division.

|  |  | Cut Detector |  |  |  |  |  |  |  |
| --- | --- | --- | --- | --- | --- | --- | --- | --- | --- |
|  |  | Normal | Midbody not detected | Tripolar divisions | Close to edge | Cut not detected | Too short cut | Missed | Total |
| Ground truth | Normal | 47 | 1 | 9 | 1 | 1 | 5 | 5 | 69 |
|  | Midbody not detected | 0 | 0 | 0 | 0 | 0 | 0 | 0 | 0 |
|  | Tripolar divisions | 2 | 0 | 4 | 0 | 0 | 0 | 1 | 7 |
|  | Close to edge | 0 | 1 | 1 | 6 | 1 | 0 | 4 | 13 |
|  | Cut not detected | 0 | 0 | 0 | 0 | 6 | 1 | 0 | 7 |
|  | Too short cut | 0 | 0 | 0 | 0 | 0 | 0 | 0 | 0 |
|  | Not a division | 0 | 1 | 1 | 0 | 0 | 1 | 0 | 3 |
|  | Total | 49 | 2 | 14 | 7 | 8 | 6 | 10 |  |

**Supplementary Table 2:** Annotated data set.

| Condition | Number of time-lapse videos |  |  |  | Number of annotated divisions |  |  |  |
| --- | --- | --- | --- | --- | --- | --- | --- | --- |
|  | Train | Val | Test | Total | Train | Val | Test | Total |
| Control | 1 | 1 | 1 | 3 | 21 | 11 | 15 | 47 |
| Control | 3 | 1 | 1 | 5 | 53 | 13 | 11 | 77 |
| CEP55 | 1 | 1 | 1 | 3 | 22 | 16 | 11 | 49 |
| Spastin | 1 | 0 | 1 | 2 | 14 | 0 | 12 | 26 |
| Total | 6 | 3 | 4 | 13 | 110 | 40 | 49 | 199 |

